# Cardiac Adaptations in the Cave Nectar Bat *Eonycteris spelaea*: Insights into Metabolic Resilience and Stress Response

**DOI:** 10.1101/2025.05.15.653669

**Authors:** Fan Yu, Akshamal M Gamage, Myu Mai Ja Kp, Randy Foo, Ying-Hsi Lin, Lijin Wang, Chee Jian Pua, Wharton Chan, Gustavo E Crespo-Avilan, Edgar M Pena, Lewis Z Hong, Aditya Iyer, Sujoy Ghosh, Elisa A Liehn, Jean-Paul Kovalik, Lin-Fa Wang, Chrishan J Ramachandra, Derek J Hausenloy

**Affiliations:** National Heart Research Institute Singapore, National Heart Centre Singapore, Singapore; Cardiovascular & Metabolic Disorders Programme, Duke-National University of Singapore Medical School, Singapore; Yong Loo Lin School of Medicine, National University Singapore, Singapore; Programme in Emerging Infectious Diseases, Duke-NUS Medical School, Singapore; Centre for Computational Biology, Duke-NUS Medical School, Singapore, Singapore; SingHealth Experimental Medicine Centre and National Large Animal Research Facility, Singapore; Paratus Sciences Singapore Pte Ltd, Singapore; Excelra, NSL Arena, Uppal, Hyderabad, India; Pennington Biomedical Research Center, Baton Rouge, Louisiana, USA; Center of Innovation and eHealth, University of Medicine and Pharmacy Carol Davila Bucharest, Romania; The Hatter Cardiovascular Institute, University College London, London, UK

**Keywords:** Cardiometabolic adaptation, Cardiac stress resilience, Comparative transcriptomics, Evolutionary cardiology, Metabolic efficiency, Bat physiology

## Abstract

**Aims:** Bats are unique mammals with remarkable adaptations, including powered flight, which demands significant energy expenditure. While previous studies have documented basic structural characteristics of bat hearts, a comprehensive understanding of their response to physiological stress remains unexplored. This study investigates the cardiac adaptations of the cave nectar bat *Eonycteris spelaea* to elucidate the mechanisms underlying their unique physiological capabilities.

**Methods and Results:** We performed RNA sequencing to analyse the cardiac gene expression profile of *E. spelaea* and 6 other bat species in comparison to mouse and human hearts, revealing enriched transcriptomic signatures related to oxidative phosphorylation and fatty acid metabolism across multiple bat species. Metabolomic profiling compared acylcarnitine metabolites and tricarboxylic acid (TCA) cycle intermediates between bat and mouse hearts, indicating a distinct acylcarnitine profile and increased levels of TCA cycle intermediates in bats, suggesting enhanced metabolic capacity. Structural adaptations were assessed through anatomical and histological analyses on cardiac tissues, showing thicker left ventricular walls and increased vascular density in bats without pathological hypertrophy. Functional characteristics were evaluated using dobutamine stress echocardiography, demonstrating superior cardiac reserve in bats with significant increases in ejection fraction, stroke volume, and cardiac output under stress conditions. Additionally, isolated cardiomyocytes were treated with Angiotensin II (Ang II) to assess stress responses. Bat cardiomyocytes displayed resistance to Ang II-induced hypertrophy and mitochondrial dysfunction compared to mice, further highlighting their resilience to stress-induced damage.

**Conclusion:** The unique adaptations observed in bat hearts, including enhanced metabolic pathways, structural remodelling, and cellular resilience contribute to their ability to meet the high energy demands of powered flight while maintain cardiac function under stress. These insights into bat cardiac physiology provide valuable information on cardioprotective mechanisms that could be applicable to other species.

**Translational Perspective:** Bats exhibit remarkable cardiac adaptations that sustain the high energy demands of powered flight while resisting stress-induced damage. These insights into evolutionarily conserved cardioprotective mechanisms highlight potential therapeutic pathways for preventing heart failure, including resistance to hypertrophy and mitochondrial dysfunction under stress. Studying non-model organisms like bats may offer innovative approaches to enhance metabolic and stress resilience in human hearts, paving the way for translational research in cardiovascular disease management and treatment.

## Introduction

Bats are a unique group of mammals characterised by several adaptations that enable powered flight and support their high metabolic demands. They exhibit an extended lifespan relative to their body size and demonstrate notable resistance to viral infections and cancer^1, 2^. As the only mammals capable of sustained flight, bats experience significantly higher energy expenditures compared to terrestrial mammals, particularly during active flight. For instance, certain frugivorous bat species can increase their basal metabolic rates by up to 15-fold and achieve heart rates as high as 900 beats per minute^3^. To accommodate these high energy requirements, bats have undergone natural selection favouring mitochondrial and nuclear genes involved in oxidative phosphorylation^4^.

The cardiovascular system of bats has evolved several distinctive features that facilitate their unique lifestyle. Bats possess the largest relative heart size among mammals, which contributes to both increased muscle mass and higher blood ejection volumes^5^. Additionally, their characteristic head-down resting position subjects the cardiovascular system to altered gravitational forces, creating sustained biomechanical stress as blood is pumped against gravity^6^. Ultrastructural and morphometric analyses have revealed significant adaptations in bat hearts. Comparative studies show that bat cardiomyocytes possess a wider T-tubule system, a higher mitochondrial volume fraction, increased crista density, and greater lipid body concentrations compared to other small mammals like hamsters and rats^7, 8^. These structural adaptations suggest enhanced metabolic efficiency and energy storage capabilities that likely contribute to their exceptional exercise tolerance.

Despite these insights into the structural characteristics of bat hearts, a comprehensive understanding of how they respond to physiological stress remains critically unexplored. Existing research has primarily focused on mitochondrial morphology and general structural features, leaving a significant gap in knowledge regarding how various bat species maintain cardiac function under sustained metabolic challenges. Given the high energy demands of flight, we hypothesise that bat hearts have evolved specific cardiometabolic adaptations that preserve myocardial energetics during prolonged stress. Our study aims to investigate the transcriptional, metabolic, structural, and functional characteristics of bat hearts at baseline conditions and under pharmacologically induced stress. By elucidating these adaptations, we aim to provide new insights into the extraordinary cardiac physiology of bats and uncover novel mechanisms of stress resilience that could inform future cardiovascular research.

## Materials and Methods

### Animal models and surgical procedures

All experiments were approved by the SingHealth Institutional Animal Care and Use Committee (IACUC) under approval numbers 2023/SHS/1847, 2015/SHS/1088, and 2020/SHS/1582, and conform to the guidelines from Directive 2010/63/EU of the European Parliament on the protection of animals used for scientific purposes. Young adult male C57BL/6J mice (10-12 weeks old) were obtained from inVivos, Jackson Laboratory, Singapore. Young adult male bats (*Eonycteris spelaea*, 18-24 months old) were sourced from a local captive-breeding colony in Singapore^9^ and housed at the National Large Animal Research Facility (NLARF).

For transverse aortic constriction (TAC) surgery, 10-week-old mice were intubated under general anaesthesia induced by intraperitoneal injection of 100 mg/kg ketamine and 10 mg/kg xylazine, with anaesthesia maintained using 2% isoflurane in oxygen. Ventilation was provided using a rodent ventilator. A left lateral thoracotomy was performed to expose the aortic arch, and the ascending aorta (between the brachiocephalic and left carotid arteries) was ligated with a silk suture using a 27G needle as a spacer, which was promptly removed after ligation. Control animals (referred to as healthy mice) underwent sham operations without aortic ligation. Mice were monitored postoperatively for behaviours changes indicative of pain or distress. Pre-operative care included analgesia (intraperitoneal injection of 0.1 mg/kg buprenorphine) and antibiotics (Baytril, 5 mg/kg in drinking water). Cardiac remodelling was assessed 8 weeks after surgery. At the terminal timepoint, mice were euthanised by intraperitoneal injection of 100 mg/kg ketamine and 10 mg/kg xylazine, while bats received an intraperitoneal injection of 100 mg/kg pentobarbital.

### RNA sequencing and transcriptomic analysis

RNA was extracted from frozen bat and mouse cardiac tissues (N=3 animals per group) using the RNeasy Plus Micro Kit (Qiagen). RNA sequencing libraries were prepared with Illumina TruSeq stranded mRNA and Nugen Ovation amplification kits, followed by sequencing on a HiSeq 4000 platform (Novogene Technology CO., Ltd, Singapore). Healthy human heart RNA-Seq data from 3 individuals were obtained from GSE116250 (SRX4297606, SRX4297613, SRX4297614). Fastq reads were aligned using the nf-core rnaseq pipeline (3.14.0) with the star_salmon workflow. Length-scaled counts were generated for each sample and differential gene expression analysis was performed using DESeq2 (padj < 0.05 and abs(logFC) > 1).

For the analysis of publicly available bat heart RNA-Seq data, fastq reads were downloaded for *Artibeus jamaicensis* (SRR13417529, SRR13417530, SRR13417531, SRR13417532), *Desmodus rotundus* (SRR13417566), *Molossus molossus* (SRR13417562), *Myotis myotis* (SRR11528219, SRR11528220), *Phyllostomus hastatus* (SRR13417565) and *Rousettus aegyptiacus* (SRR2913354, SRR2914359). Since raw fastq data was not readily available for human GTEx left ventricle (LV) samples, publicly deposited TPM data for human GTEx (V8) was downloaded for analysis. A complete TPM matrix included 432 human GTEx LV samples, 3 *E. spelaea* heart samples, 3 mouse heart samples, and 11 publicly available bat samples from 6 different species. 15,230 genes that were annotated in all samples were used for differential gene expression analysis using Wilcoxon tests^10^ and significant DEGs were identified using the criterion: abs(logFC) > 1 and FDR < 0.01. Pathway analyses of significant DEGs were performed against the MSigDB hallmark and GO-BP gene sets using the genekitr ORA (over-representation test) function^11^.

### Metabolomic profiling of acylcarnitines and organic acids

Acylcarnitine and organic acid (TCA cycle intermediates) metabolic profiling analyses were performed at the Duke-NUS Metabolomics Facility. Cardiac tissues were weighed and homogenised in ice-cold 50% acetonitrile containing 0.3% formic acid at a concentration of 50 mg tissue per mL of homogenate. Organic acids were extracted from the tissue homogenate and derivatised to their trimethylsilyl forms before quantitation by gas chromatography-mass spectrometry (GC-MS). Acylcarnitines and amino acids were extracted and derivatised to their methyl and butyl esters, respectively, then quantitated using liquid chromatography-mass spectrometry (LC-MS). The results were normalised to tissue weight to account for variations in sample input.

### Western blotting

Western blots were performed using a previously described protocol^12^. Proteins (25 μg) extracted from animal cardiac tissues were separated on 4-12% Bis-Tris Bolt Gels (Thermo Fisher Scientific, MA, USA) and transferred to nitrocellulose (NC) membranes using the iBlot Dry Blotting System (Thermo Fisher Scientific). NC membranes were blocked and incubated overnight with primary antibodies: CD36 (1:500; #ab64014, Abcam, Cambridge, UK) and GLUT4 (1:1000; #9001, Cell Signaling Technology, MA, USA). The following day, membranes were washed, incubated with HRP-conjugated secondary antibodies, and developed using SignalFire™ ECL Reagent (Cell Signaling Technology). Tropomyosin (1:20,000, #3910, Cell Signaling Technology) was used as a loading control. Blots were imaged using the C-DiGit Blot Scanner (LI-COR, NE, USA) and densitometry was performed using Image Studio software (LI-COR).

### Transmission electron microscopy

Left ventricular tissues from mice and bats were processed for transmission electron microscopy (TEM) using a standardised protocol. Tissue fragments were fixed in 0.1M phosphate buffer (2% paraformaldehyde, 2% glutaraldehyde) at 4°C, washed, and post-fixed in 2% osmium tetroxide with potassium ferrocyanide for 2 hours at room temperature. After dehydration through an ethanol series, tissues were embedded in resin and ultrathin sectioned at 90nm using an EM UC7 ultramicrotome (Leica Microsystems, Wetzlar, Germany). Ultrastructural imaging was performed on a FEI Tecnai Spirit G2 TEM (FEI Company, OR, USA) with an FEI Eagle 4k digital camera at 2,000x magnification.

### Histological and immunofluorescence analysis

Cardiac tissues were harvested, washed with ice-cold phosphate-buffered saline, and immediately fixed in 10% neutral buffered formalin. After processing and paraffin embedding, tissues were sectioned at 4 µm thickness (Leica AG, Solms, Germany). Tissue sections were deparaffinised using Histo-Clear and rehydrated through a graded ethanol series (100% to 70%). For morphological analysis, Gömöri trichrome staining was performed. Fibrosis was assessed using a Sirius Red and Fast Green kit (#9046, Chondrex, WA, USA). Images were acquired using a Zeiss Axiovert 200M microscope (Carl Zeiss, Oberkochen, Germany), and the fibrotic area was quantified as a percentage of the total image area using ImageJ software, with five sections analysed per sample.

For immunofluorescence, antigen retrieval was performed using Bull’s Eye Solution (#BULL1000MX, Biocare Medical, CA, USA) at 98°C for 10 minutes. Tissue sections were blocked with 5% BSA for 45 minutes, then incubated with WGA Alexa Fluor™ 488 Conjugate (1:200, #W11261, Thermo Fisher Scientific, MA, USA) for 3 hours at room temperature. Tissues were mounted using TrueVIEW with DAPI (#SP-8500-15, Vector Laboratories, CA, USA). Images for cardiomyocyte cross-sectional area analysis were acquired using a fluorescence microscope (Carl Zeiss), with approximately 100 cells measured per sample. For CD36 and GLUT4 immunostaining, antigen retrieval was performed using citrate buffer (#ab64214, Abcam, Cambridge, UK) at 98°C for 10 minutes. Tissue sections were blocked with 5% BSA for 1 hour and incubated overnight with primary antibodies: CD36 (1:200; #ab64014, Abcam), GLUT4 (1:200; 1:200; #ab48547, Abcam), cardiac Troponin T (1:200, #ab10214, Abcam), and α-actinin (1:200; #ab137346, Abcam). The following day, sections were washed and incubated with Alexa Fluor^TM^ 488 (#A-11070) and Alexa Fluor^TM^ 555 (#A-21422) secondary antibodies (both from Thermo Fisher Scientific). Tissues were mounted using TrueVIEW with DAPI (#SP-8500-15, Vector Laboratories).

To measure isolated cardiomyocyte size, cells were fixed with 4% paraformaldehyde, permeabilised with 0.3% Triton X-100, blocked with 5% BSA, and incubated overnight with anti-sarcomeric α-actinin primary antibodies (#ab137346, Abcam). Cells were then probed with Alexa Fluor™ Plus 488 secondary antibodies (#A-11070, Thermo Fisher Scientific) and counterstained with DAPI. Representative images of isolated cardiomyocytes were captured using an inverted confocal microscope (LSM710, Carl Zeiss). All image analyses were performed using ImageJ software.

### Echocardiography

Echocardiography was performed using a Vevo 2100 system (Visual Sonics Inc., Toronto, Canada) equipped with a 38 MHz MicroScan transducer (frequency range 18-38 MHz) at the SingHealth Experimental Medicine Centre (SEMC). Anaesthesia was induced with 5% isoflurane and maintained at 1% isoflurane throughout the procedure. A comprehensive examination included two-dimensional (2D) B-mode imaging, motion-mode (M-mode) imaging, and tissue Doppler imaging (TDI). Images were acquired at baseline and following an intraperitoneal bolus injection of dobutamine (10 μg/g body weight). The same anaesthesia protocol was maintained throughout the procedure to ensure comparability between baseline and post-dobutamine measurements.

### Myofibril mechanics assay

Myofibril mechanics were quantified as previously described^13^. Briefly, the fast solution switching technique was used to assess myofibril contractile properties. Left ventricular (LV) sections were skinned overnight in rigor solution (132 mM NaCl, 5 mM KCl, 1 mM MgCl_2_, 10 mM Tris, 5 mM EGTA, pH 7.1) containing 0.5% Triton X-100, protease inhibitors (10 μM leupeptin, 5 μM pepstatin, 200 μM PMSF, and 10 μM E64), 500 μM NaN_3_, and 500 μM DTT at 4°C. Skinned LVs were homogenised in relaxing solution (pCa 9.0, where pCa = -log10[calcium concentration]). Myofibril suspensions were transferred to a temperature-controlled chamber (15°C) and mounted between two microtools: one connected to a motor for length changes (Mad City Labs), the other a calibrated cantilevered force probe (12.2 μm/μN; frequency response 2-5 kHz). Myofibril length was set at approximately 2.2 μm. Sarcomere lengths and myofibril diameters were measured using ImageJ software (NIH). Myofibrils were activated and relaxed by rapid translation between solutions of different pCa. Mechanical and kinetic parameters measured included resting tension, maximal tension, rate constant of tension development (kACT), duration of linear relaxation, and rate constant of exponential relaxation (fast kREL). At least one sample from each group was analysed daily, with all cultured cells in an experiment harvested on the same day. While experiments were not blinded during conduct, data analysis was performed in a blinded fashion.

### Cardiomyocyte isolation and culture

Cardiomyocytes from mice and bats were isolated using a Langendorff-free direct needle perfusion method^14^. Animals were anaesthetised with 75% CO_2_/25% O_2_ at 1 L/min flow rate containing isoflurane, maintained at 2% via nose cone. The heart was flushed by injecting EDTA buffer into the right ventricle, then excised and transferred to fresh EDTA buffer. Cardiac tissues were digested with collagenase buffer until softened. Ventricles were separated, and the tissue was gently dissociated by pipetting. The cell suspension was filtered through a 100-µm filter, and cardiomyocytes were collected by gravity sedimentation for 20 minutes, followed by three additional sedimentations in calcium reintroduction buffers. Yield and viability were assessed using a haemocytometer. For culture, cardiomyocytes were suspended in M199 medium (#M4530, Merck, Darmstadt, Germany) supplemented with 5% FBS, 10 mmol/L BDM (2,3-butanedione monoxime), and Penicillin-Streptomycin, then plated on laminin-coated (15 µg/mL) dishes for 1 hour at 37°C in 5% CO_2_. Media was then changed to culture media with or without 10 μM Angiotensin II (#ab120183, Abcam, Cambridge, UK) for 24 hours. Culture media comprised M199 media supplemented with 0.1% BSA, 1% Insulin-Transferrin-Selenium (#I3146, Merck, Darmstadt, Germany), 10 mM BDM, 1% CD lipid (#11905, Thermo Fisher Scientific), and Penicillin-Streptomycin.

### Mitochondrial respiration assay

Cardiomyocytes were seeded at 4×10³ cells/well on Seahorse 96-well XF Cell Culture Microplates (Agilent Technologies, CA, USA) pre-coated with laminin (15 µg/mL). Prior to the assay, culture media was replaced with Seahorse XF Media supplemented with 10 mM glucose, 2 mM glutamate and 1 mM sodium pyruvate, and mitochondrial function was assessed using the Seahorse XF Cell Mito Stress Test Kit (Agilent Technologies). Compounds were sequentially injected at the following final concentrations: oligomycin (2.5 μM), FCCP (1 μM), and a mixture of antimycin A and rotenone (5 μM each). Mitochondrial respiration parameters were calculated as previously described^15^.

### Statistical analysis

Statistical analysis was performed using GraphPad Prism 10.1.2, with data expressed as mean ± SEM. Normality was assessed using Shapiro-Wilk test (n<8) or D’Agostino and Pearson test (n≥8). For normally distributed data, two-tailed unpaired Welch’s t-tests were used for two-group comparisons, while one-way ANOVA with Tukey post hoc test was applied for multiple groups. Non-normally distributed data were analysed using Mann-Whitney U tests for two groups or Kruskal-Wallis tests with Benjamini-Hochberg false discovery rate correction for multiple groups. A *P*-value of <0.05 was considered statistically significant.

## Results

### Bat hearts exhibit enriched transcriptomic signatures related to energy production and metabolism

We investigated the cardiac gene expression profile of the cave nectar bat *Eonycteris spelaea* by conducting RNA sequencing analysis and comparing it with mouse and human hearts. Principal component analysis revealed distinct clustering of samples from the three species, with a notable observation that human hearts exhibit greater similarity to bat hearts than to mouse hearts (Figure 1A). Differential gene expression analysis uncovered 11,449 differentially expressed genes (DEGs) between bats and humans (7,042 downregulated and 4,407 upregulated), and 8,185 DEGs between bats and mice (3,907 downregulated and 4,278 upregulated) (Figure 1B). A pairwise study of enriched biological pathways demonstrated that bat hearts uniquely exhibited significant enrichment in oxidative phosphorylation, fatty acid metabolism, and adipogenesis pathways. In contrast, human hearts showed enrichment in epithelial-mesenchymal transition, myogenesis, and apical junction pathways (Figure 1C).

**Figure 1:**
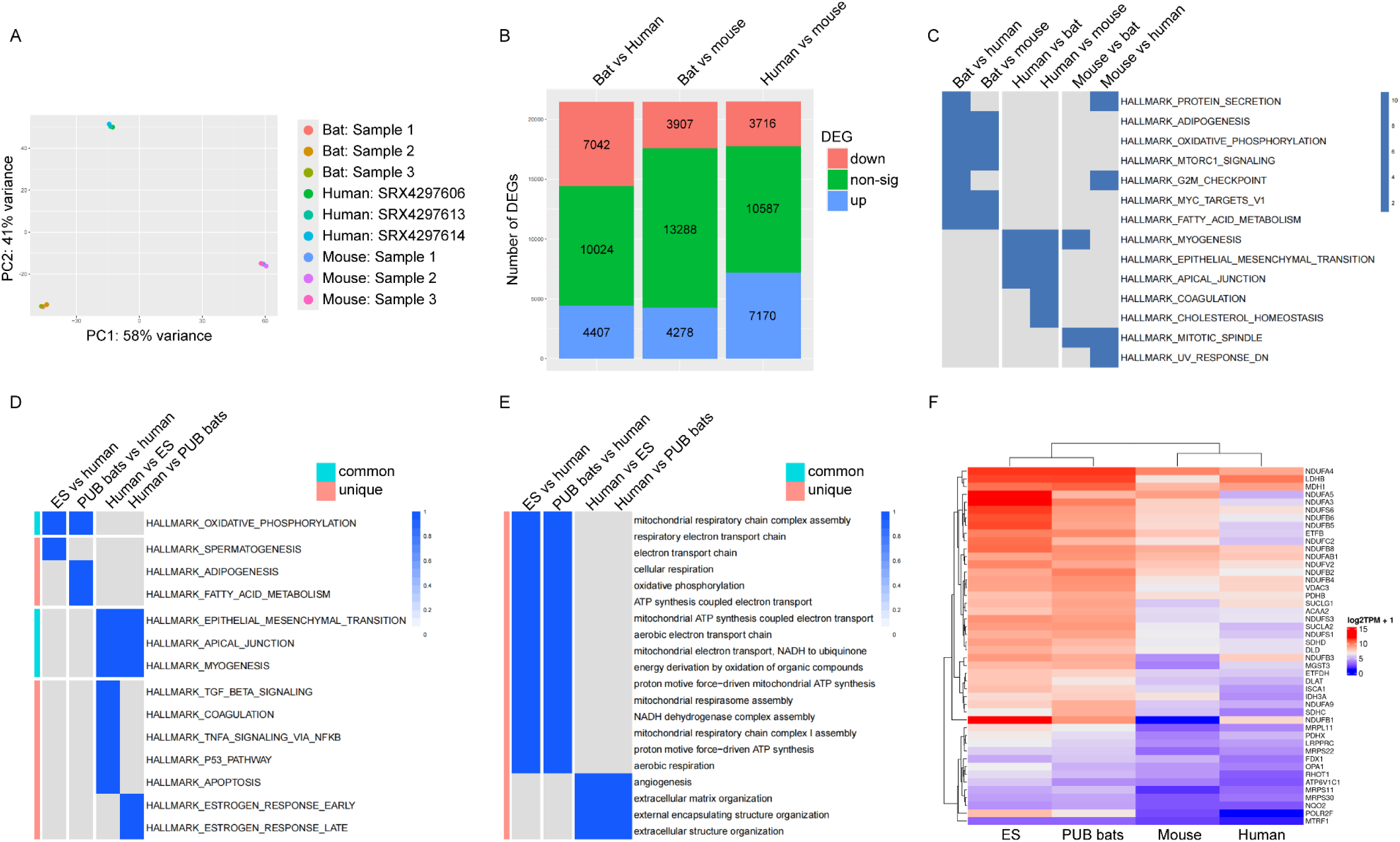
Comparative transcriptomics and pathway analysis of bat, human, and mouse cardiac tissues. (A) Principal component analysis plot showing clustering of cardiac tissue samples by species. (B) Quantification of differentially expressed genes (DEGs) among *E. spelaea*, human, and mouse cardiac tissues, highlighting inter-species transcriptomic differences. (C) Comparison of enriched hallmark pathways in *E. spelaea*, human, and mouse cardiac tissues, showing species-specific and shared cellular processes. (D) Comparison of enriched hallmark pathways across *E. spelaea* (ES), 6 other bat species (PUB bats), and human cardiac tissue, showing evolutionary conserved metabolic processes in bats. (E) Comparison of enriched GO-BP pathways across 7 bat species and human cardiac tissue, showing evolutionary conserved mitochondria- and energy-related processes in bats. (F) Heatmaps of DEGs in the oxidative phosphorylation hallmark pathway across 7 bat species, human, and mouse cardiac tissue, showing evolutionary conserved upregulation of electron transport chain genes in bats. For panels C-E, significant pathways in the heatmaps are represented by blue tiles, as indicated by - log10(padj). In panels D and E, common pathways refer to those enriched in more than one comparison, while unique pathways are those enriched in only one comparison. Abbreviations: PUB-publicly available.

To determine the broader applicability of these metabolic pathway adaptations, we analysed heart RNA-seq data from an additional 6 bat species (11 samples in total; Table 1) and compared them with 432 human left ventricle (LV) heart samples from the GTEx database. Pathway analysis using the MSigDB hallmark gene set consistently revealed upregulation of oxidative phosphorylation, fatty acid metabolism, and adipogenesis pathways across all 7 bat species compared to humans (Figure 1D). Gene Ontology Biological Process (GO-BP) analysis further corroborated these findings, demonstrating significant enrichment for mitochondria- and energy-related biological processes in bat hearts (Figure 1E). A heatmap derived from the oxidative phosphorylation hallmark pathway illustrated the upregulation of genes predominantly involved in the electron transport chain across all bat species relative to humans and mice (Figure 1F). Collectively, these findings reveal a conserved metabolic signature across diverse bat species, suggesting fundamental evolutionary adaptations in cardiac energy metabolism. These adaptations, likely driven by natural positive selection^4^, may underpin the extraordinary physiological capabilities observed in bats. From this point forward, the data presented is from *E. spelaea* only and will be referred to as "bats".

**Table 1:** Ecological and behavioural traits of bats used in this study.

| Species | Diet | Geography | Hibernation | Migration |
| --- | --- | --- | --- | --- |
| *Eonycteris spelaea* | Nectar | South / Southeast Asia | No | No |
| *Artibeus jamaicensis* | Fruit | Central / South America | No | No |
| *Desmodus rotundus* | Blood | Central / South America | No | No |
| *Molossus molossus* | Insect | Central / South America | No | No |
| *Myotis myotis* | Insect | Europe | Yes | No |
| *Phyllostomus hastatus* | Generalist | Central / South America | No | No |
| *Rousettus aegyptiacus* | Fruit | Africa, Middle East | No | Yes |

### Bat hearts exhibit enhanced metabolic capacity and unique metabolic adaptations

To elucidate the adaptive metabolomic profile in bats, we analysed acylcarnitine metabolites, key indicators of fuel selection in cardiac metabolism^16–18^. The metabolomics results were consistent with our transcriptomic findings, which revealed enrichment in pathways related to fatty acid metabolism and oxidative phosphorylation. Serum samples revealed a distinctive acylcarnitine profile in bats, with increased concentrations of short-(C4, C5), medium-(C6, C8, C10), and long-chain acylcarnitine (C14, C16, C18:1) compared to mice (Figure 2A and C). Cardiac tissues exhibited alterations in the short-chain acylcarnitine profile, with elevated levels of C3, C4, and C5 (metabolic derivatives of amino acids and fatty acids) in comparison to mice (Figure 2B and D). Notably, long-chain acylcarnitines (C14, C16, C18) were also elevated in bat hearts, although these differences did not reach statistical significance. Additionally, bat hearts showed increased levels of tricarboxylic acid (TCA) cycle intermediates including pyruvate, succinate, fumarate, and malate (Figure 2E), suggesting an expanded TCA intermediate pool to support enhanced oxidative phosphorylation and exercise capacity^19^.

**Figure 2:**
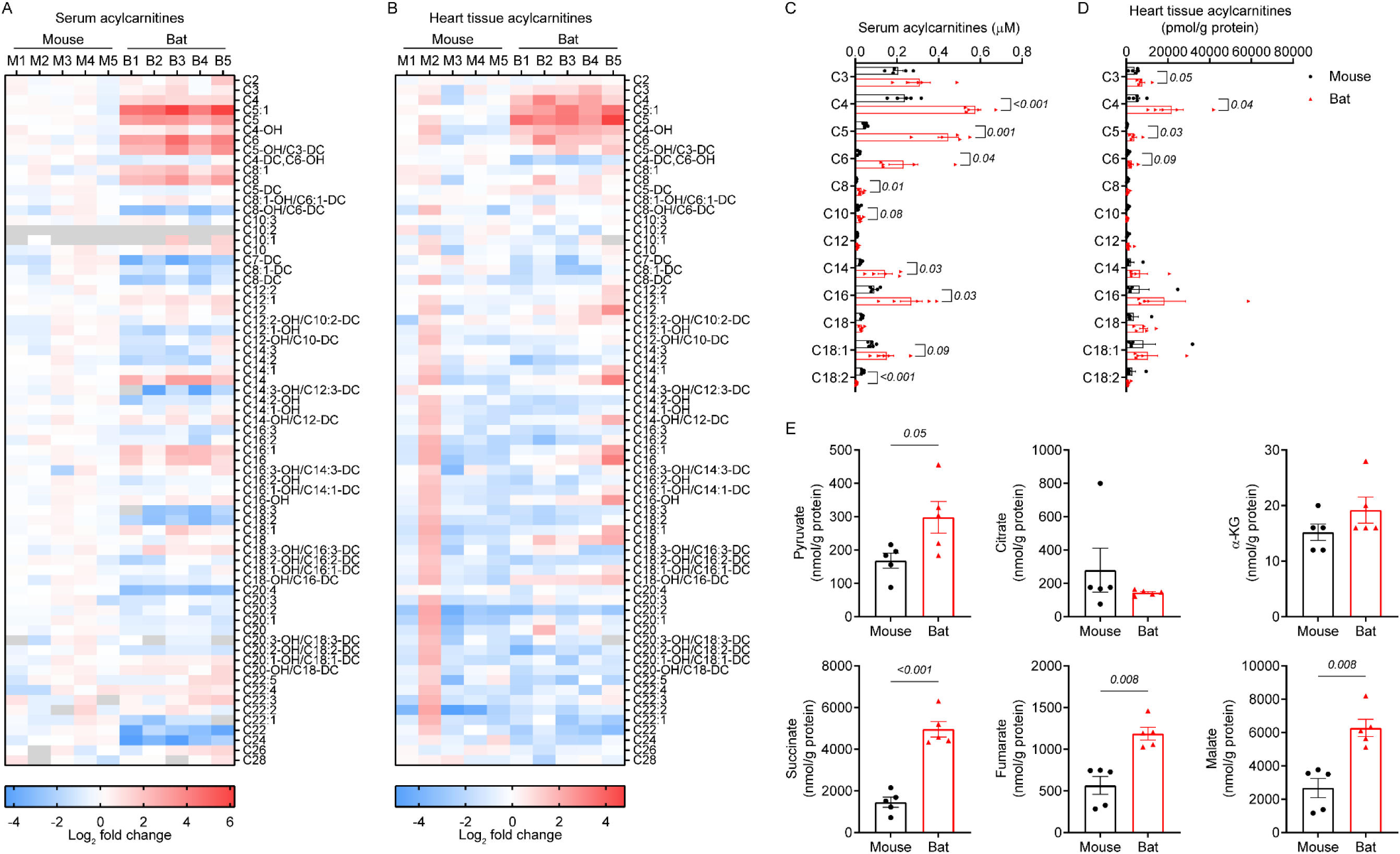
Comparative metabolomic analysis of serum and cardiac tissues in bats and mice. (A) Heatmap of serum acylcarnitine profiles. (B) Heatmap of cardiac tissue acylcarnitine profiles. (C) Selected serum acylcarnitines showing increased levels of short-, medium, and long-chain acylcarnitines in bats compared to mice. (D) Selected cardiac tissue acylcarnitines showing increased levels of mostly short-chain acylcarnitines in bats compared to mice. (E) TCA cycle intermediates showing increased levels of pyruvate, succinate, fumarate, and malate in bat hearts compared to mouse hearts. Data are presented as mean ± SEM (N=5 animals per group). Statistical analysis: Welch’s t-test (panels C, D, and E for pyruvate and succinate); Mann-Whitney U test (panel E for fumarate and malate).

Given that bat flight muscles support both carbohydrate and fatty acid oxidation^20, 21^, we examined the expression of critical substrate transporters in cardiac tissue. Immunohistochemical staining revealed that both GLUT4 (glucose transporter type 4) and CD36 (cluster of differentiation 36) were expressed in cardiac tissues of bats and mice (Figure 3A and B, Supplementary figure 1A and 1B). However, consistent with the upregulation of metabolic pathways observed in our transcriptomic analysis, western blot analysis confirmed that bat hearts expressed higher levels of both GLUT4 and CD36 compared to mouse hearts (Figure 3C, Supplementary figure 2), indicative of better metabolic flexibility. Collectively, these findings reveal a distinctive cardiac fuel utilisation strategy in bats, characterised by enhanced metabolic capacity and a distinct cardiometabolic profile. Such adaptations likely enable bats to efficiently meet the fluctuating energy demands associated with powered flight and their unique ecological niches^20,22, 23^.

**Figure 3:**
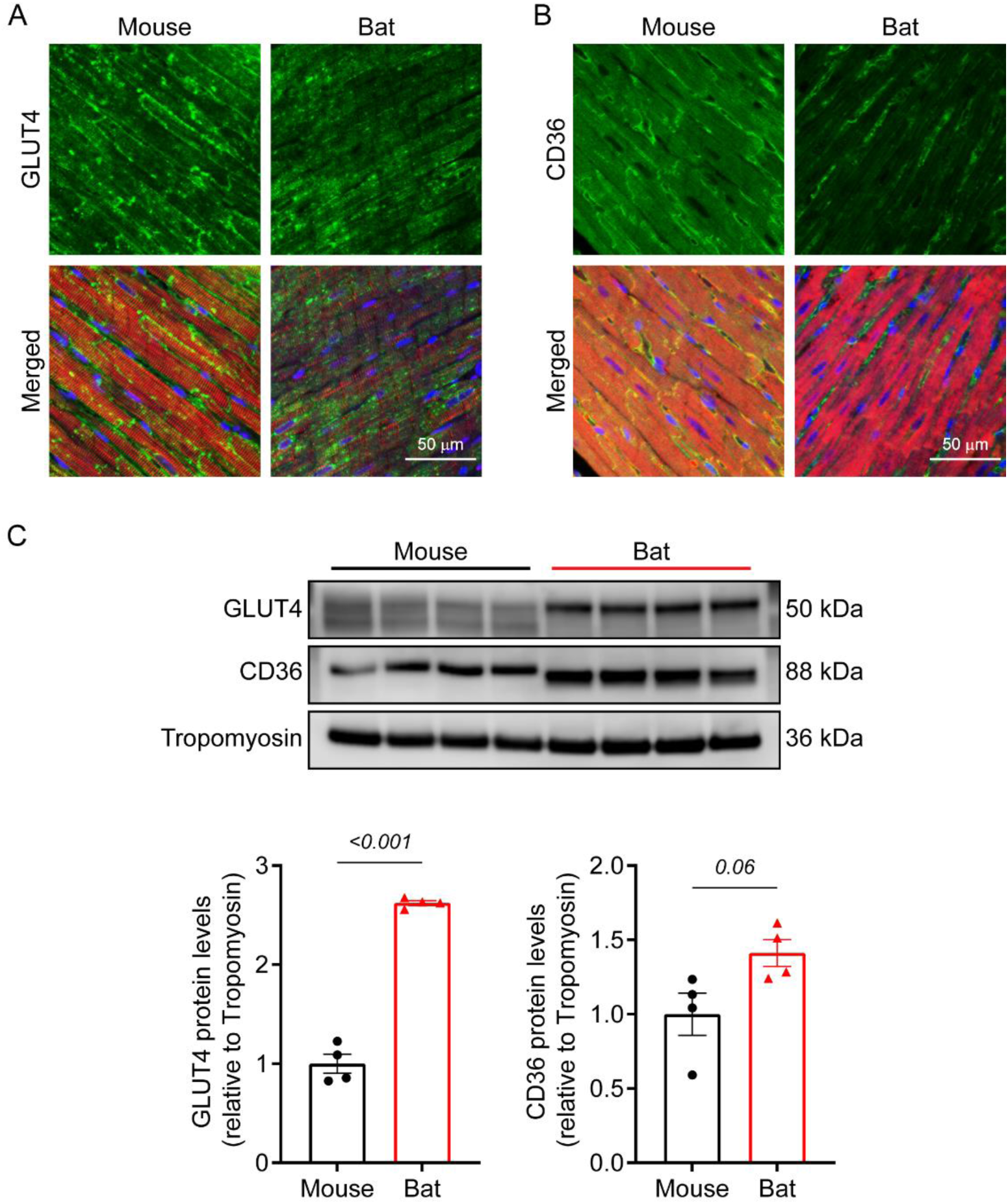
Expression of metabolic transporters in bat and mouse hearts. (A) Immunofluorescence images showing the expression of GLUT4 in bat and mouse cardiac tissue, counterstained with α-actinin (red) and DAPI (blue). (B) Immunofluorescence images showing the expression of CD36 in bat and mouse cardiac tissue, counterstained with cardiac troponin T (cTnT; red) and DAPI (blue). (C) Western blot analysis showing increased protein levels of GLUT4 and CD36 in bat hearts compared to mouse hearts. Data are presented as mean ± SEM (N=4 animals per group; Welch’s t-test).

### Bat hearts exhibit signs of adaptive remodelling and stress

Building upon the observed transcriptomic and metabolic adaptations, we next investigated whether these unique features were accompanied by structural modifications in bat hearts compared to mouse hearts. Detailed anatomical analysis revealed striking differences between bat and mouse hearts, with bat hearts showing notable similarities to human hearts. Longitudinal bisection from apex to base showed that bat hearts possess well-developed atrial chambers, similar to those observed in humans^24^, in contrast to the rudimentary atrial chambers observed in mice (Figure 4A). Moreover, bat ventricles exhibited a more elongated shape, akin to human hearts, compared to the ellipsoidal shape of mouse ventricles.

**Figure 4:**
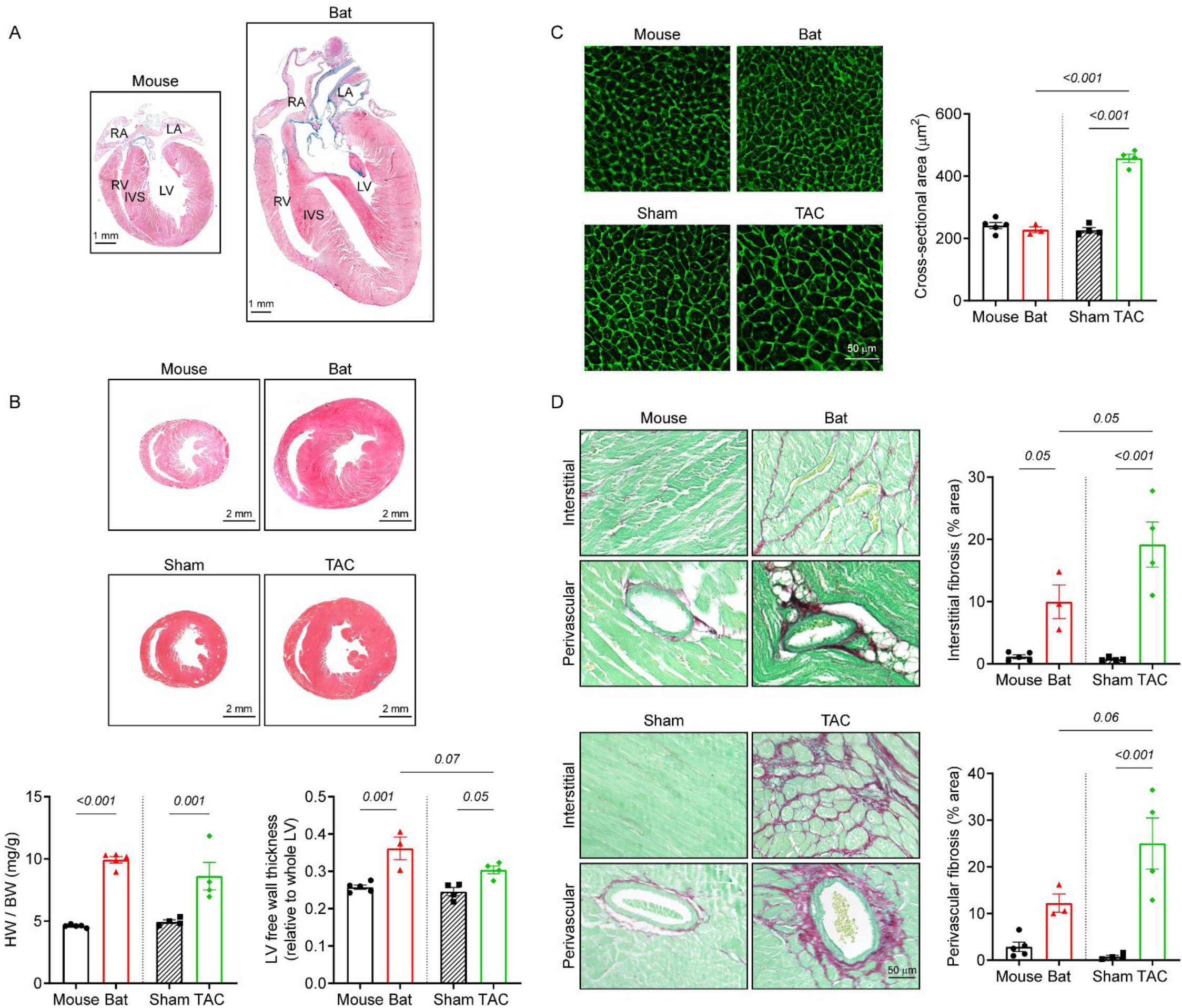
Structural comparison of adult bat, healthy mouse, and TAC mouse hearts. (A) Anatomical comparison of mouse and bat hearts. (B) Cardiac tissue cross-sections stained with Gömöri trichrome, showing that bats and TAC mice have larger hearts with thickened LV walls compared to healthy mice. (C) Cardiac tissue sections stained with wheat germ agglutinin, showing cardiomyocyte hypertrophy exclusively in the TAC mouse heart. (D) Cardiac tissue sections stained with Sirius Red/Fast Green, showing increased interstitial and perivascular fibrosis in TAC mouse hearts and to a lesser extent in bat hearts. Data are presented as mean ± SEM (N=3-5 animals per group; One-way ANOVA with Tukey post hoc test). Abbreviations: RA- right atrium; LA- left atrium; RV- right ventricle; LV- left ventricle; IVS- interventricular septum.

Quantitative analysis demonstrated that bat hearts have a significantly larger relative size compared to healthy mouse hearts (bat vs mouse; 9.9 ± 0.3 mg/g vs 4.6 ± 0.1 mg/g), and this increased heart weight was comparable to that of hypertrophied mice induced by transverse aortic constriction (TAC) (8.6 ± 1.1 mg/g). Assessment of cardiac remodelling revealed that bat hearts had strikingly thicker LV free walls than healthy mouse hearts, with the increase in LV thickness slightly exceeding that of TAC mice (Figure 4B). Interestingly, despite the increased heart size and wall thickness, we observed no significant difference in cardiomyocyte size between bats and healthy mice (Figure 4C), suggesting the absence of pathological cellular hypertrophy in bats^25^. In contrast, TAC mice exhibited cellular hypertrophy, as evidenced by enlarged cardiomyocyte size (Figure 4C). Further examination of cardiac fibrosis revealed that bat hearts displayed more perivascular and interstitial fibrosis compared to healthy mouse hearts (Figure 4D). In contrast, TAC mouse hearts showed substantial amounts of both perivascular and interstitial fibrosis (Figure 4D). These observations indicate an adaptive response in bat hearts potentially induced by biomechanical stress.

Transmission electron microscopy analysis revealed more densely packed mitochondrial networks in bat hearts compared to mouse hearts (Figure 5A). Additionally, we observed perivascular adipocytes located near microvasculature and larger vessels in bat hearts, which were absent in mouse hearts (Figure 5B). This observation may explain the enrichment of adipogenesis pathways in bats identified in our transcriptomics analysis. Quantification of vascular density showed a greater number of vessels per area in bat hearts compared to mouse hearts (bat vs mouse; 5.3 ± 0.3 per mm^2^ vs 1.6 ± 0.5 per mm^2^; Figure 5C). Collectively, these structural modifications suggest a sophisticated cardiac remodelling mechanism that differs fundamentally from traditional mammalian cardiac physiology. These adaptations likely contribute to the enhanced cardiac function and stress tolerance observed in bats, potentially supporting their unique physiological demands, particularly in meeting the high energy requirements of powered flight.

**Figure 5:**
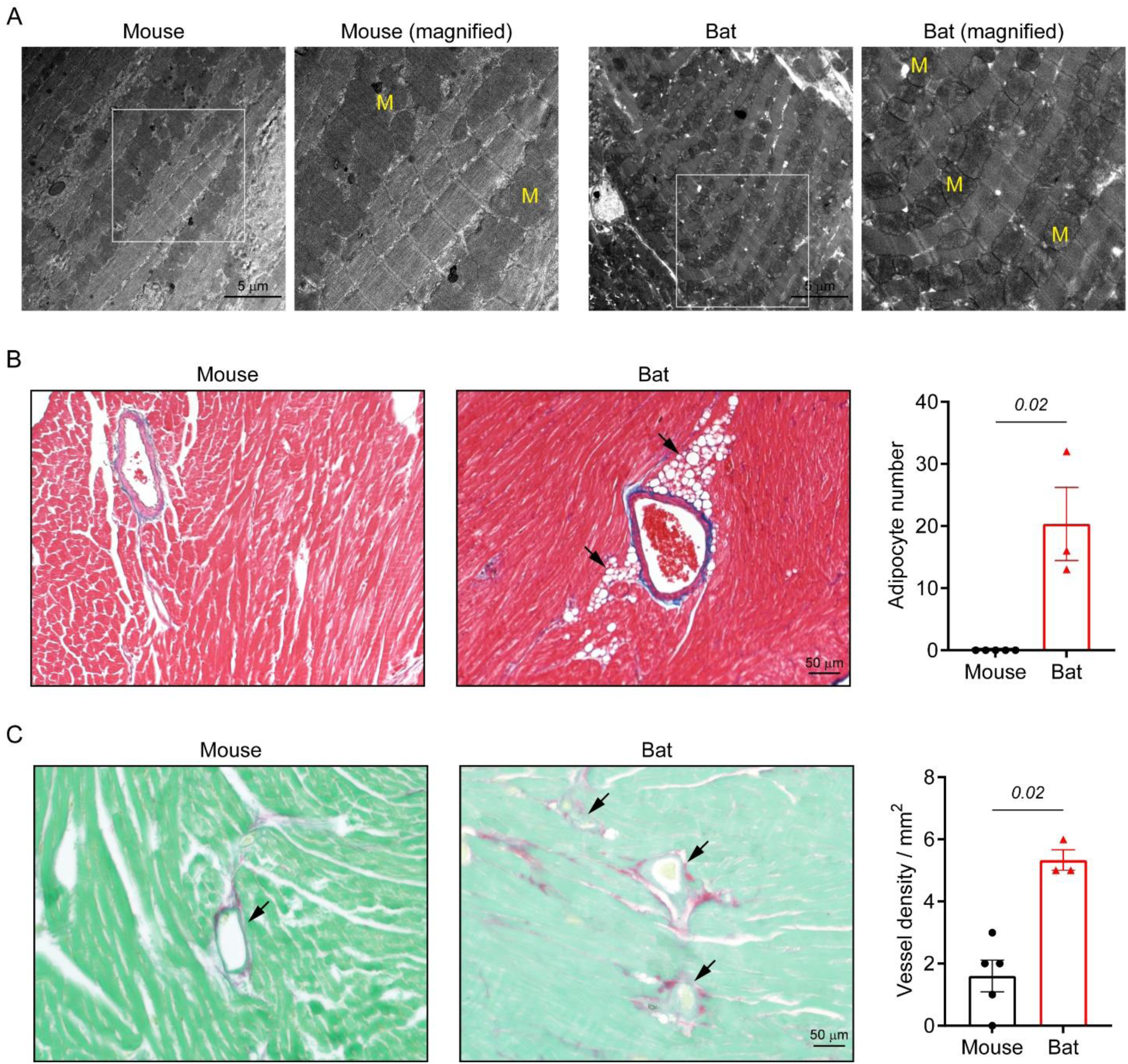
Ultrastructural and vascular characteristics of bat and mouse hearts. (A) Transmission electron microscopy images showing denser mitochondrial networks in bat hearts compared to mouse hearts. White insets represent magnified regions with ‘M’ denoting mitochondria. (B) Cardiac tissue sections showing perivascular adipocytes (black arrows) adjacent to vessels in bat hearts, a feature absent in mouse cardiac tissue. (C) Cardiac tissue sections stained with Sirius Red/Fast Green, showing higher vascular density (black arrows) in bat hearts compared to mouse hearts. Data are presented as mean ± SEM (N=3-5 animals per group; Mann-Whitney U test).

### Bat hearts reveal superior cardiac reserve and stress responsiveness

We next investigated the functional characteristics of bat hearts under baseline and stressed conditions. At baseline, heart rates in anesthetised bats were comparable to those in anesthetised mice when positioned supine and subjected to the same isoflurane concentration. Consistent with our earlier structural findings, bats demonstrated larger LV mass compared to mice (Table 2). Despite their larger body surface area due to wing size, stroke volume and cardiac output remained similar between species. Notably, ejection fraction (EF) and fractional shortening (FS) were significantly lower in bats than in mice, suggesting distinct cardiac functional profiles at baseline (Table 2).

**Table 2:** Cardiac function of bats and mice at baseline and after dobutamine administration.

|  | Baseline |  |  | Dobutamine |  |  |
| --- | --- | --- | --- | --- | --- | --- |
|  | Mouse | Bat | *P*-value | Mouse | Bat | *P*-value |
| Heart rate, beats/min | 464.3 ± 12.4 | 450.8 ± 19.5 | ns | 493.0 ± 8.8 | 487.0 ± 25.7 | ns |
| Left ventricular mass, mg | 123.2 ± 4.4 | 433.1 ± 30.9 | 0.0018 | 134.0 ± 13.2 | 392.5 ± 53.6 | 0.0142 |
| Left ventricular ejection fraction, % | 62.8 ± 1.9 | 30.0 ± 4.7 | 0.0031 | 83.1 ± 1.2 | 66.1 ± 3.2 | 0.0081 |
| Fractional shortening, % | 40.0 ± 5.1 | 17.7 ± 6.2 | 0.057 | 59.2 ± 2.3 | 48.3 ± 1.7 | 0.0103 |
| Relative wall thickness | 0.5 ± 0.02 | 0.6 ± 0.1 | ns | 0.6 ± 0.0 | 0.9 ± 0.0 | 0.0024 |
| Stroke volume, µL | 46.8 ± 1.9 | 59.7 ± 6.7 | ns | 56.6 ± 2.8 | 99.6 ± 9.1 | 0.0139 |
| Cardiac output, mL/min | 21.8 ± 1.4 | 27.3 ± 4.3 | ns | 27.9 ± 1.6 | 49.1 ± 7.1 | 0.0555 |

Data are presented as means ± SEM (N=4 animals per species). Statistical significance is indicated by *P*-values; ns indicates not significant.

To determine whether the reduced baseline cardiac function represented a potential limitation or an adaptive characteristic, we conducted dobutamine stress echocardiography. A single bolus of dobutamine (10 µg/g) was administered intraperitoneally to both species to evaluate their cardiac response^26^. After dobutamine injection, EF and FS remained lower in bats compared to mice, while relative wall thickness, stroke volume, and cardiac output significantly increased in bats (Table 2). Representative M-mode echocardiograms confirmed our previous structural observations, revealing larger LV wall thicknesses in bats and greater baseline contractility in mice (Figure 6A). Interestingly, both species demonstrated similar chronotropic responses, with comparable increases in heart rates following dobutamine administration (Figure 6B). However, the inotropic response in bat hearts was substantially more pronounced. Bat hearts exhibited a remarkable increase in EF compared to mouse hearts (bat vs mouse; 135.8 ± 35.2% vs 32.93 ± 5.3%; Figure 6B). The percentage change in FS was approximately 4-fold greater in bats versus mice (bat vs mouse; 255.1 ± 82.6% vs 57.4 ± 25.7%; Figure 6B). Moreover, stroke volume and cardiac output in bats increased nearly 3-fold, far exceeding the response observed in mice (Figure 6B). Bat hearts also demonstrated a notable increase in relative wall thickness compared to mouse hearts, although this difference did not reach statistical significance (Figure 6B).

**Figure 6:**
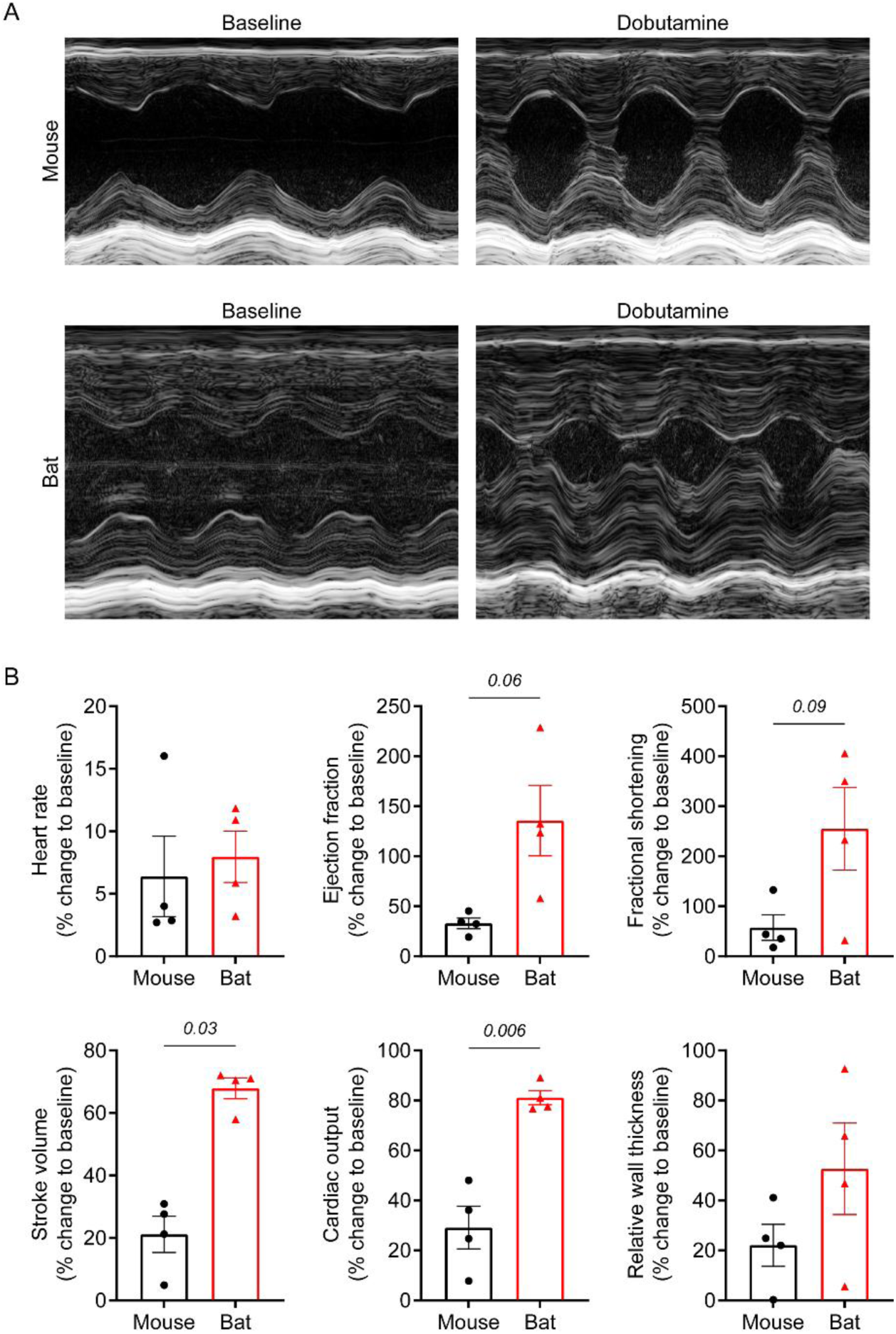
Comparative cardiac response to dobutamine in bats and mice. (A) Representative M-mode echocardiography images of bat and mouse hearts at baseline and after dobutamine injection. (B) Effects of dobutamine on heart rate, ejection fraction, fractional shortening, stroke volume, cardiac output, and relative wall thickness in bats and mice. Data are presented as mean ± SEM (n=4 animals per groups). Statistical analysis: Welch’s t-test (panel B for ejection fraction, fractional shortening, and cardiac output); Mann-Whitney U test (panel B for stroke volume)

To further elucidate the mechanisms underlying this superior inotropic response, we compared bat cardiac myofibrils with mouse cardiac myofibrils. Although force generation and activation kinetics were similar between the two species (Figure 7A and B), bats showed a non-significant tendency towards faster linear phase relaxation, with a slight decrease in duration and increase in rate of relaxation (Figure 7C and D). Most notably, bat myofibrils demonstrated approximately 2-fold faster exponential phase relaxation kinetics (bat vs mouse; 36.1 ± 2.0 s^-1^ vs 17.4 ± 2.1 s^-1^; Figure 7E). This enhanced relaxation may support quicker transitions between contraction and relaxation states, particularly contributing to improved diastolic function and enabling more effective filling of the heart between contractions^27^. These findings align with our previous metabolic and structural observations, suggesting that unique adaptations in bat hearts, including enhanced mitochondrial networks, metabolic capacity, and structural remodelling contribute to their exceptional cardiac reserve.

**Figure 7:**
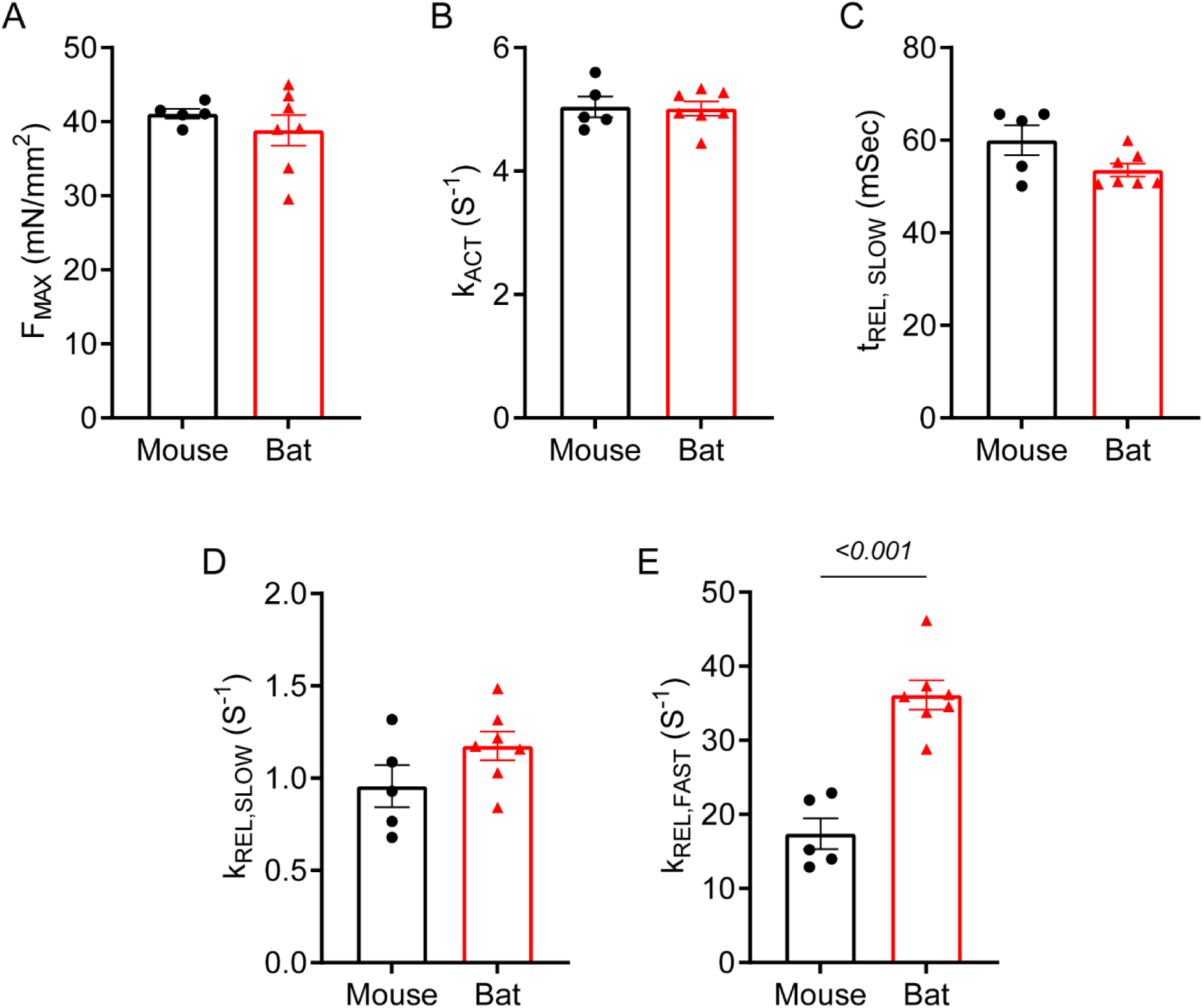
Myofibril mechanics in bat and mouse cardiomyocytes. (A) Force generation in bat and mouse myofibrils. (B) Activation kinetics, represented by the rate constant of tension development (kACT), in bat and mouse myofibrils. (C) Duration of linear phase relaxation in bat and mouse myofibrils. (D) Linear phase relaxation kinetics in bat and mouse myofibrils. (E) Exponential phase relaxation kinetics, represented by the rate constant of exponential relaxation (fast kREL), in bat and mouse myofibrils. Data are presented as mean ± SEM (N=5-7 animals per group; Welch’s t-test for panel E).

### Bat cardiomyocytes demonstrate unique stress response

Given the low basal cardiac function yet increased cardiac reserve in bats, we speculated that this is the result of cardiometabolic adaptations that allow for the preservation of cardiac energetics during conditions of pathophysiological stress. Angiotensin II (Ang II), as the effector molecule of the renin-angiotensin system, has been shown to induce cardiomyocyte hypertrophy and mitochondrial dysfunction in neonatal rat cardiomyocytes^28, 29^. To evaluate whether bat cardiomyocytes have evolved with cardiometabolic adaptations, isolated ventricular cardiomyocytes from adult bats and mice were treated with Ang II (10 µM) for 24 hours, followed by assessment of cell size and mitochondrial function.

While Ang II treatment elicited cellular hypertrophy in isolated mouse cardiomyocytes, it failed to induce hypertrophy in bat cardiomyocytes (Figure 8A and B). Additionally, Ang II mediated mitochondrial dysfunction in mouse cardiomyocytes, as evidenced by reduced basal respiration, maximal respiration, and spare reserve capacity (Figure 8C). However, these impairments were not found in bat cardiomyocytes (Figure 8D). These findings suggest that bat cardiomyocytes possess unique adaptations that confer resistance to Ang II-induced hypertrophy and mitochondrial dysfunction. This resistance may be linked to the previously observed metabolic and structural characteristics of bat hearts, indicating a robust capability to maintain cardiac function under stress conditions.

**Figure 8:**
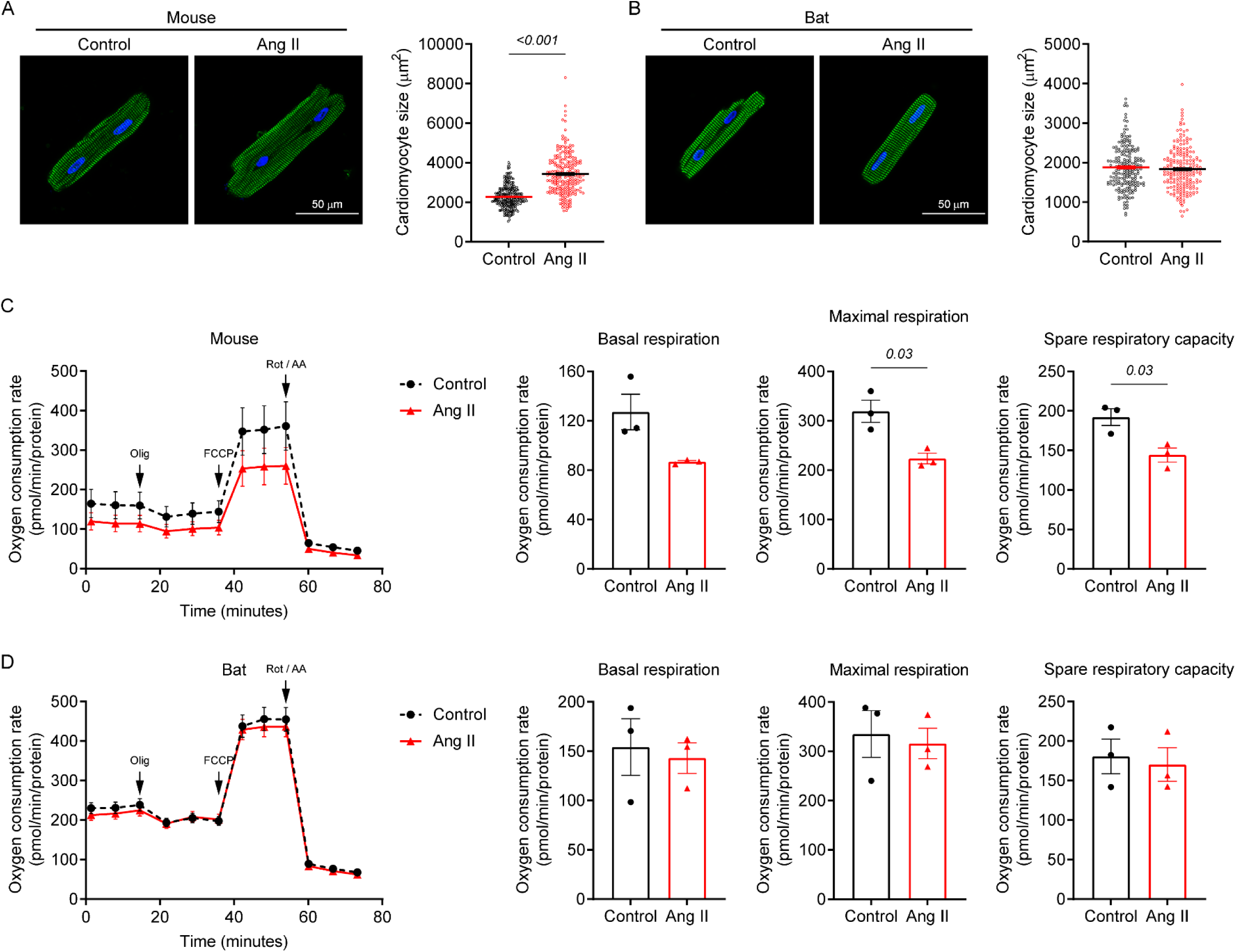
Differential effects of Angiotensin II on cardiomyocyte hypertrophy and mitochondrial function in bats and mice. (A) Immunofluorescence images showing Ang II induced cellular hypertrophy in mouse cardiomyocytes. (B) Immunofluorescence images showing no effect of Ang II on bat cardiomyocyte size. (C) Oxygen consumption rate trace of mouse cardiomyocytes showing Ang II-induced mitochondrial functional impairment (left) and changes in respiratory parameters (right) (D) Oxygen consumption rate trace of bat cardiomyocytes showing no effect of Ang II on mitochondrial function (left) and no changes in respiratory parameters (right). Data are presented as mean ± SEM (N=4 animals per group, 50-73 cells per animal for cellular hypertrophy assessment; N=3 animals per group for mitochondrial function analysis). Statistical analysis: Mann-Whitney U test (panels A and B); Welch’s t-test (panel C for maximal respiration and spare respiratory capacity). Abbreviations: Olig-oligomycin; Rot-rotenone; AA-antimycin A.

## Discussion

Our comprehensive investigation of bat cardiac adaptations reveals several novel findings. (1) Transcriptomic analysis across 7 bat species demonstrated enriched signatures in oxidative phosphorylation and fatty acid metabolism, suggesting a conserved evolutionary metabolic strategy. (2) Metabolomic profiling indicated a distinct acylcarnitine profile with elevated TCA cycle intermediates, revealing enhanced metabolic capacity. (3) Functional and structural analyses showed superior cardiac reserve, characterised by thicker LV walls, increased vascular density, significant stress-induced performance improvements, and remarkable cardiomyocyte resistance to hypertrophic and mitochondrial stress. These findings collectively highlight the extraordinary physiological adaptations that enable bat hearts to meet the extreme energy demands of powered flight.

Our findings on the cardiac structure of *E. spelaea* corroborate and extend previous studies on bat cardiac adaptations. *E. spelaea* exhibits larger hearts relative to body size compared to mice, which are non-flying mammals, supporting higher specific oxygen uptake during flight^30^. The increased vascular density in *E. spelaea* hearts align with observations of higher capillary density in bat cardiac and skeletal muscles, particularly in smaller species^31^. Ultrastructural analysis revealed denser mitochondrial networks in *E. spelaea* cardiomyocytes, consistent with findings in other bat species. Studies on *Eidolon helvum* and *Pipistrellus pipistrellus* have shown higher proportions of mitochondria in bat cardiomyocytes compared to terrestrial mammals, with unique arrangements suggesting adaptations for efficient energy utilisation^7, 8^. These observations collectively indicate evolutionary specialisation for enhanced aerobic metabolism and increased energy production capacity in bat cardiac tissue. Novel findings include the presence of perivascular adipocytes and increased cardiac fibrosis. The latter may result from cumulative ’wear and tear’ due to prolonged inversion during roosting, as previous studies have shown such behaviour can cause cardiac damage in bats^6^. However, the observed fibrosis may serve an adaptive function, providing structural support for hearts subjected to extreme physiological demands during both roosting and flight. Despite increased heart size and wall thickness, we observed no significant cardiomyocyte hypertrophy, indicating a form of physiological cardiac remodelling distinct from pathological hypertrophy^25^. These structural adaptations collectively contribute to the remarkable cardiac performance of bats, enabling them to meet the extreme physiological demands of powered flight.

Bats, as the only mammals capable of powered flight, have evolved remarkable metabolic adaptations to meet their high energy demands. Our transcriptomic analysis of seven bat species revealed significant enrichment of pathways related to fatty acid metabolism and oxidative phosphorylation in bat hearts compared to human and mouse hearts. This consistent upregulation across species suggests an evolutionary adaptation through positive selection, indicating a primary reliance on fatty acid utilisation in bat hearts. Our metabolomic findings in *E. spelaea* further support this conclusion, showing increased levels of short-, medium-, and long-chain acylcarnitines in serum, and elevated short-chain acylcarnitines and TCA cycle intermediates in cardiac tissue. This profile suggests enhanced fatty acid β-oxidation and increased oxidative phosphorylation in bat hearts. The discrepancy between serum and cardiac tissue acylcarnitine profiles likely reflects the dynamic nature of acylcarnitine metabolism, with serum levels indicating overall metabolic state and tissue levels representing local metabolic processes. The accumulation of short-chain acylcarnitines in *E. spelaea* cardiac tissue may be attributed to the bats’ nocturnal feeding habits and potential fasting state during tissue harvesting, aligning with previous studies on hibernating and aroused bats^32^. These metabolic adaptations in bat hearts are particularly intriguing given the diverse dietary and physiological challenges faced by bats across different ecological niches. Despite limited research on cardiac metabolic flexibility in bats, evidence shows they rapidly and efficiently utilise ingested nutrients. Nectarivorous species use dietary sugar as an immediate energy supply during flight, while insectivorous bats primarily oxidise their protein-rich diet^22^. Studies on the Egyptian fruit bat *Rousettus aegyptiacus* and nectar bat *Glossophaga soricina* demonstrate that bats can efficiently use recently ingested nutrients for both resting and flight metabolism, with up to 80% of hovering flight energy coming from recently ingested sugar in some species^20, 22, 23^. While immediate energy utilisation pathways may be species-dependent, bats likely possess conserved pathways to provide energy during fasting periods, as they must survive on daily food intake and face a constant risk of starvation^3^. We speculate that while dietary energy provides immediate fuel, fatty acid metabolism likely serves as the main energy source during periods of fasting or prolonged flight. A notable observation in our study was the presence of perivascular adipocytes, coupled with enrichment of adipogenesis pathways across the seven bat species studied. This suggests potential energy reserves, supported by previous findings of significant adipose mass in bat hearts^3, 5, 33^. These fat deposits likely serve as a crucial energy reservoir during active flying and food scarcity, with frugivorous bats capable of replacing up to half of their fat reserves within a single day of foraging^3^. Our findings also revealed significantly increased expression of GLUT4 and CD36 transporter proteins in *E. spelaea* cardiac tissue, likely allowing for metabolic flexibility during hover feeding and fasting. These adaptations collectively enhance the metabolic flexibility and capacity of bat hearts, potentially contributing to their extraordinary flight capabilities.

The functional analysis of *E. spelaea* hearts revealed novel insights into bat cardiac physiology. At baseline, *E. spelaea* exhibited significantly lower ejection fraction (EF) and fractional shortening (FS) compared to mice, aligning with the unique cardiac innervation system observed in other bat species^34^. This reduced basal function likely reflects an evolutionary adaptation for energy conservation during rest. However, upon dobutamine stimulation, *E. spelaea* hearts demonstrated a remarkable increase in EF, FS, stroke volume, and cardiac output, far exceeding the response in mice. These findings parallel observations in other species adapted to extreme physiological demands, such as the cyclic bradycardic state in *Uroderma bilobatum*^3^ and the cardiac plasticity of naked mole-rats^35^. This enhanced cardiac reserve could suggest adaptations in β-adrenergic sensitivity, calcium handling, and metabolic flexibility. Our myofibril data, showing approximately 2-fold faster exponential phase relaxation kinetics in *E. spelaea* cardiomyocytes, may reflect superior calcium handling similar to that observed in the superfast muscles of echolocating bats, where rapid calcium transients and increased fibre shortening velocities are critical for extreme physiological performance^36^. The superior cardiac reserve observed in bat hearts, coupled with their enhanced myofibrillar relaxation kinetics, provides compelling evidence of an evolved mechanism for meeting extreme metabolic demands. The exceptional cardiac adaptability of *E. spelaea* likely underpins bats’ capacity to meet the metabolic demands of powered flight and apparent resistance to cardiovascular diseases. These findings suggest a remarkable cardiac plasticity that expands our understanding of cardiac performance and adaptation in mammals, offering valuable insights for human cardiac research.

Our investigation into the response of *E. spelaea* cardiomyocytes to Angiotensin II (Ang II) revealed a remarkable resistance to pathological alterations typically associated with cellular hypertrophy and mitochondrial dysfunction^28^. This resistance is particularly noteworthy given that Ang II has been previously demonstrated to induce cardiac hypertrophy through increased mitochondrial reactive oxygen species (ROS) production. Two potential explanations for this phenomenon were considered. Initially, we hypothesised that the observed resistance might be due to incompatibility between the human-origin Ang II ligand used in our culture system and the bat angiotensin II type 1 receptor (AT1R). However, protein sequence analysis revealed functionally conserved AT1R domains across species, with bat-human and bat-mouse comparisons both displaying 97% similarity (comparable to the 98% similarity observed in mouse-human AT1R), effectively ruling out ligand-receptor incompatibility. This evolutionary conservation effectively rules out ligand-receptor incompatibility as an explanatory factor. The more plausible explanation lies in the unique downstream mechanisms present in bat cardiomyocytes. This hypothesis is supported by previous studies on bat physiology and longevity. For instance, research on the little brown bat *Myotis lucifugus* demonstrated that their mitochondria produce significantly less hydrogen peroxide per unit of oxygen consumed compared to shorter-lived non-flying mammals, supporting the free radical theory of aging^37^. Furthermore, studies on South American bat species have revealed exceptionally high antioxidant defences, including elevated superoxide dismutase and catalase activities, as well as remarkably high levels of α-tocopherol and β-carotene in various tissues^38^. These enhanced antioxidant mechanisms, coupled with their ability to modulate antioxidant defences during daily torpor-activity transitions, likely contribute to bats’ unique ecophysiological adaptations and extended lifespan^38^. Collectively, these findings suggest that the resistance of *E. spelaea* cardiomyocytes to Ang II-induced pathological changes is likely due to evolved mechanisms for managing oxidative stress and maintaining cellular homeostasis. This adaptation may play a crucial role in bats’ ability to withstand the high metabolic demands of flight while simultaneously contributing to their exceptional longevity. These results reveal a novel cellular stress resistance mechanism with potential implications for understanding cardiac resilience across species. This unique cardiomyocyte physiology provides a new perspective on evolved cardioprotective mechanisms and inform future research on cardiac stress responses in other species.

Our study, while providing novel insights into bat cardiac adaptations, is constrained by several key limitations. The research primarily focused on *E. spelaea* and a limited number of bat species, potentially restricting the generalisability of our findings across bat diversity. Mechanistic uncertainties persist regarding the precise molecular pathways underlying cardiomyocytes’ stress resistance, and our *in vitro* experiments may not fully capture the complex physiological conditions of active flight. These limitations underscore the need for broader, more comprehensive studies to fully elucidate the extraordinary cardiac adaptations that enable bats’ remarkable physiological capabilities.

## Conclusion

Our investigation reveals that bat hearts exhibit unique metabolic, structural, and functional adaptations that support their physiological performance during powered flight. These adaptations include enhanced metabolic capacity, superior cardiac reserve, and notable resistance to stress. By highlighting the sophisticated cardiac biology of bats, this study provides insights into metabolic efficiency and resilience that contributes to our understanding of cardioprotection and evolutionary biology.

## Acknowledgements

The authors thank Nicole Tee from the SingHealth Experimental Medicine Centre (SEMC) for echocardiography acquisition and analysis.

## Supplementary Data

**Supplementary figure 1:**
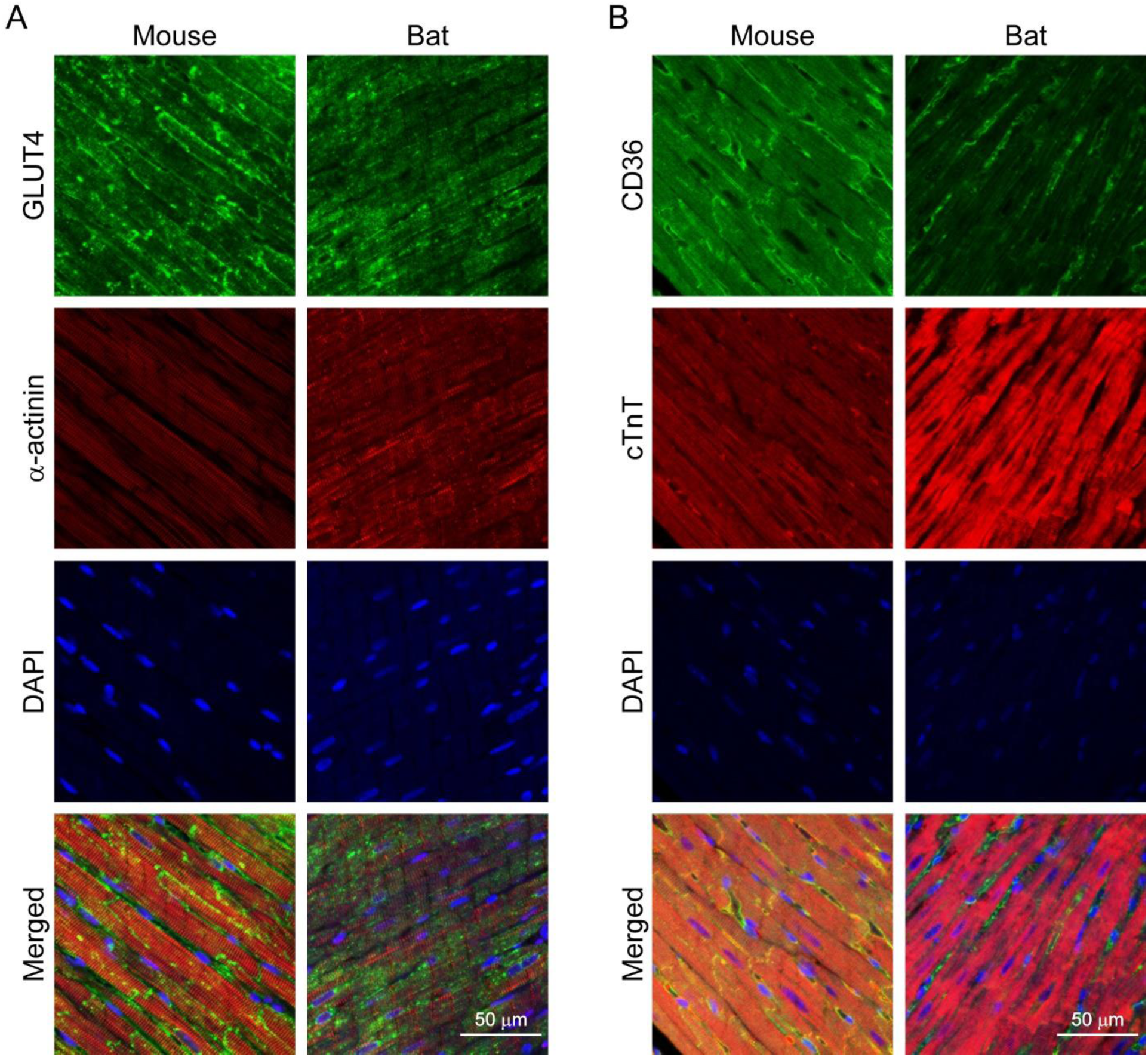
(A) Single-channel and merged immunofluorescence images of bat and mouse cardiac tissue stained with GLUT4, α-actinin, and DAPI. (B) Single-channel and merged immunofluorescence images of bat and mouse cardiac tissue stained with CD36, cardiac troponin T (cTnT), and DAPI.

**Supplementary figure 2:**
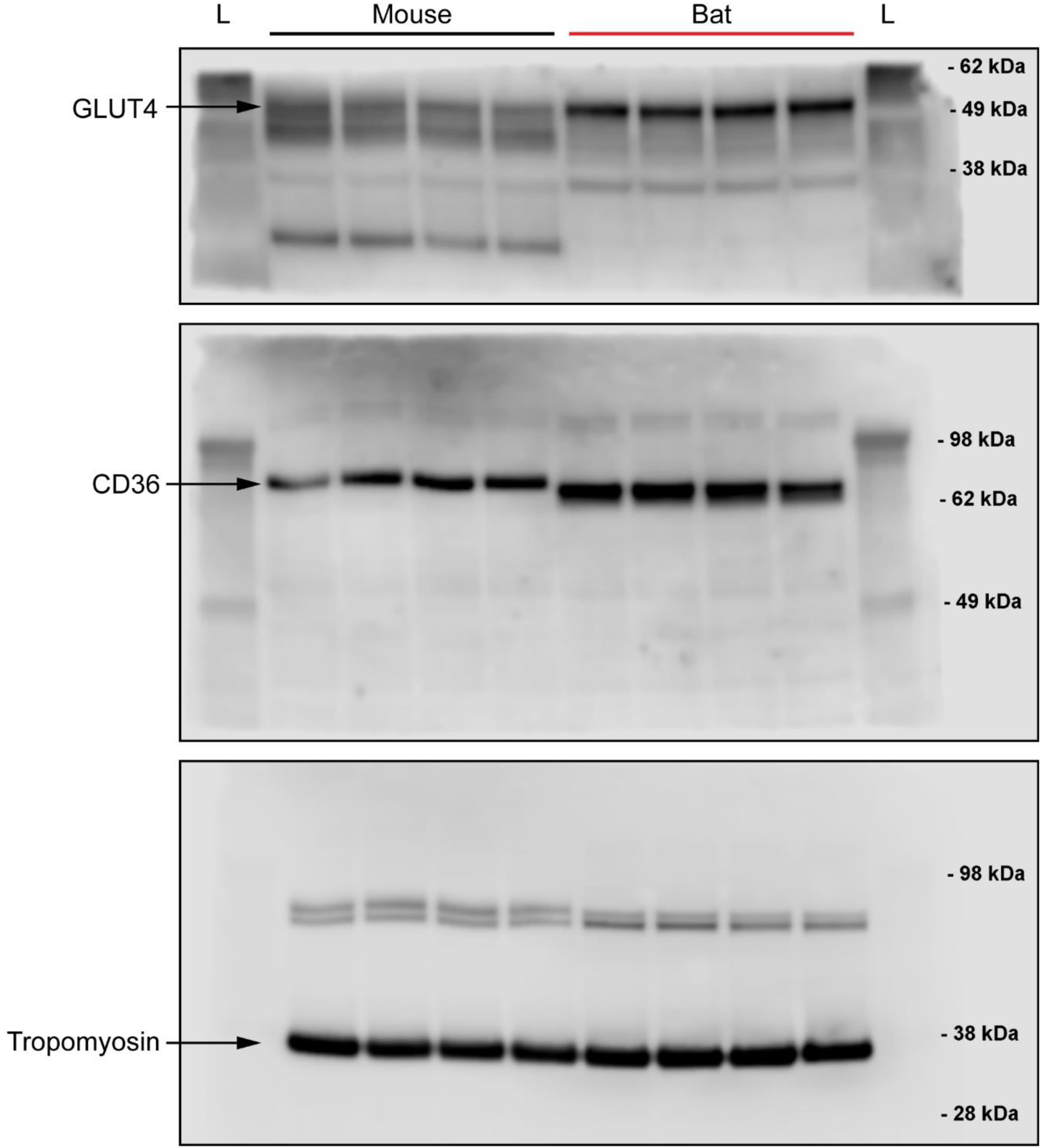
Uncropped immunoblots for detecting GLUT4 and CD36 protein levels in bat and mouse cardiac tissue. Tropomyosin was used as a loading control. Abbreviations: L-molecular weight ladder.

